# Reversible multiscale corneal sexual dimorphism orchestrates neuro-epithelial coupling and tear proteome composition

**DOI:** 10.1101/2025.10.02.680047

**Authors:** Nadege Feret, Alicia Caballero Megido, Sarah Pernot, Melissa Girard, Marilou Decoudu, Laura Fichter, Aurore Attina, Jerome Vialaret, Alexandre David, Vincent Daien, Karine Loulier, Christophe Hirtz, Frederic Michon

**Author notes:** These authors contributed equally to the study (Nadege Feret, Alicia Caballero Megido, Sarah Pernot).

## Abstract

Donor–recipient sex mismatch is an under-recognized risk factor for corneal graft rejection, yet the biological underpinnings at the ocular surface remain undefined. We hypothesized that sex hormones coordinately tune the tear–nerve–epithelium axis and that androgen exposure could normalize sex-linked differences relevant to alloimmune risk. We profiled 12-week-old mice in three hormonal contexts, i.e. male, female, and testosterone-treated female, combining tear biochemistry and label-free proteomics with epithelial lineage dynamics, in vivo sensory function, and corneal/trigeminal transcriptional readouts. Tear collection rate was comparable across groups, but total tear protein was reduced in females. Proteomics revealed extensive sex differences in extracellular composition, spanning lipid transport, protease–antiprotease balance, complement activity, and secretory/mucin pathways. Epithelial analyses showed sex-linked differences in progenitor output and spatial deployment, and sensory metrics indicated divergent innervation architecture and function. Across modalities, androgen supplementation in females shifted molecular and physiological profiles toward the male state, attenuating nearly all dimorphic signals, demonstrating that close to all sexual dimorphism observed here is reversible by testosterone and reflects an actively maintained endocrine state rather than a fixed developmental program. Complementary metabolite profiling provides a sex-stratified atlas of free modified nucleosides in tears, positioning extracellular epitranscriptomic markers as accessible indicators of hormonal context. Together, these results establish a multiscale, hormone-responsive sexual dimorphism at the ocular surface and offer a mechanistic framework linking sex and endocrine status to parameters that influence graft integration. They motivate sex-aware biomarkers, stratification by hormonal context in diagnostics and trials, and therapeutic strategies, including androgenic modulation, to mitigate the elevated rejection risk associated with sex mismatch in corneal transplantation.

## Introduction

Sexual dimorphism, the phenomenon by which members of different sexes of the same species display distinct anatomical or physiological features, has long been a subject of interest in fields spanning evolutionary biology (Delph, 2003), ecology (Shine, 1989), and genetics (Fairbairn & Roff, 2006). While it is often illustrated by obvious differences in reproductive organs or secondary sexual characteristics, such as plumage in birds (Roulin & Jensen, 2015) or antler size in deer (Malo *et a*l, 2005), it also manifests in subtler ways throughout the body. An increasing body of evidence demonstrates that numerous organ systems, from the cardiovascular system (Lock *et al*, 2021) to the nervous system (Joel *et al*, 2018), can be influenced by sex-based differences in gene expression, hormonal signaling, and developmental processes, leading to sexual dimorphism in organ physiology. Recognizing and understanding these variations is crucial not only from a theoretical standpoint but also for developing personalized and inclusive approaches to healthcare (Stolarz & Rusch, 2015). By systematically examining how sex-linked biological factors shape organ function, clinicians and researchers can adapt their practice to the physiology of patients according to their specific needs. By systematically examining how sex-linked biological factors shape organ function, clinicians and researchers can adapt their practice to the physiology of patients according to their specific needs. This study aims to fill this gap by dissecting how hormonal context shapes the corneal microenvironment across molecular, cellular, and neuro-functional levels.

The eye, and the cornea in particular, is hormone-responsive: androgen, estrogen, and progesterone receptors are present in ocular surface tissues and adnexa, and experimental and clinical data link sex steroids to ocular surface homeostasis, meibomian lipid synthesis, wound healing, and immune tone (Sullivan *et al*, 2000, 2017; Schirra *et al*, 2006). Yet an integrated, systems-level analysis of hormone-dependent corneal phenotypes remains limited.

Focusing on the cornea is justified biologically and clinically. Biologically, the cornea is the most densely innervated tissue, enabling tight neuro⍰epithelial coupling that governs epithelial renewal, tear reflexes, and barrier integrity; it is also unusually accessible to non⍰invasive imaging (*in vivo* confocal microscopy) and to topical interventions, facilitating translation. Clinically, common corneal disorders show sex⍰linked patterns: dry eye disease is more prevalent in women (Sullivan *et al*, 2017), and linked to androgen signaling in the meibomian gland (Schirra *et al*, 2006); keratoconus shows sex⍰associated differences in several cohorts; and some transplant studies implicate donor–recipient sex mismatch in graft outcomes—together pointing to actionable sexual dimorphism at the ocular surface (Debourdeau *et al*, 2022).

Bridging to the corneal tear–nerve–epithelium axis, peer⍰reviewed evidence supports direct hormonal influence. Androgens up⍰regulate lipogenic programs in the meibomian gland, promoting tear film stability (Schirra *et al*, 2006); sex steroids are detectable in human tears by LC⍰MS/MS, providing a proximate route for local signaling (Robciuc *et al*, 2022); and *in vivo* work demonstrates sex⍰ and estradiol⍰dependent differences in corneal nerve regeneration and innervation metrics. mechanisms that could reshape corneal sensitivity, epithelial turnover, and trophic support in a sex⍰dependent manner (Pham *et al*, 2019).

The existing literature relevant to the cornea has catalogued sex⍰linked signals but largely interrogates single layers (glandular, epithelial, or neural) rather than the integrated system that maintains corneal clarity. Consensus reports and focused reviews emphasize sex hormones as key determinants of ocular surface disease (Sullivan *et al*, 2017; Kelly *et al*, 2023), yet a comprehensive, multi⍰omic and functional analysis of hormone context across the corneal system is still needed as ophthalmological treatments have been applied to patients, regardless of their sex. By honing our understanding of how these differences play out in each organ, including those that appear superficially identical between sexes, we can work toward eliminating potential gaps in treatment protocols. We hypothesize that the sex hormone context (male, female, and testosterone⍰supplemented female) has a direct impact on the molecular landscape of the tear proteome, corneal transcriptome, and trigeminal neurone signature, Furthermore, we expect the modification of sex hormone levels to transform this landscape, as well as the function of these elements, yielding distinct sexual dimorphism at tissue and systems levels. By resolving these mechanisms, our study aims to provide a mechanistic basis for sex⍰aware corneal therapeutics, with implications for dry eye management and optimization of donor–recipient matching strategies in transplantation.

Here, we set out to determine whether circulating androgens coordinately shape the corneal tear– nerve–epithelium axis and whether these effects are reversible. We profiled mice in three hormonal contexts, i.e. male, female, and testosterone-treated female, to quantify tear film chemistry and proteomes, map epithelial renewal dynamics, and assess sensory architecture and function alongside corneal and trigeminal transcriptomes. This integrated design allowed us to test, within the same animals, (i) whether sex associates with distinct extracellular molecular signatures in tears, (ii) whether epithelial progenitor output and its spatial deployment differ by sex and hormonal state, and (iii) whether corneal innervation and upstream ganglionic programs covary with endocrine context. In the sections that follow, we report a multiscale sexual dimorphism across these modalities that is reversed by androgen supplementation, thereby linking hormone status to extracellular composition, neural organization and sensitivity, and epithelial renewal logic. These results provide a framework for sex-aware biomarkers and interventions at the ocular surface and motivate stratification by hormonal context in both discovery and clinical translation.

## Results

### Sexual hormones govern specific aspects of the tear film parameters

Corneal homeostasis relies on a finely tuned crosstalk among the tear film, dense sensory innervation, and epithelial paracrine signaling. Disruption of any one of these axes compromises barrier integrity and can precipitate sight-threatening disease; for example, tear deficiency in dry eye can drive epithelial thickening, haze, and neovascularization (Sacchetti *et al*, 2020).

Extracellular modified ribonucleosides—particularly methylated and pseudouridylated species—are emerging as sensitive, integrative readouts of cellular RNA metabolism (McCown *et al*, 2020). Because they enter biofluids only after turnover of their parent RNAs, each circulating modification reflects the coupled dynamics of RNA synthesis, editing, and decay, as well as the activity of the enzymes that install or erase these marks. Perturbations in these pathways have been linked to cancer, metabolic disease, and mitochondrial dysfunction (Boughanem *et al*, 2023).

Against this backdrop, we asked whether the ocular surface bears a sex-biased nucleoside signature. By profiling 26 modified ribonucleosides in basal tear film, we uncovered pronounced dimorphism: eight (31%) were enriched in females (**Fig. 1A**) and six (23%) in males (**Fig. 1B**), such that more than half (14/26; 54%) differed by sex. Among the twelve which did not exhibit a sexual dimorphism, most of them were methylations (**Fig. 1C**).

**Figure 1.**
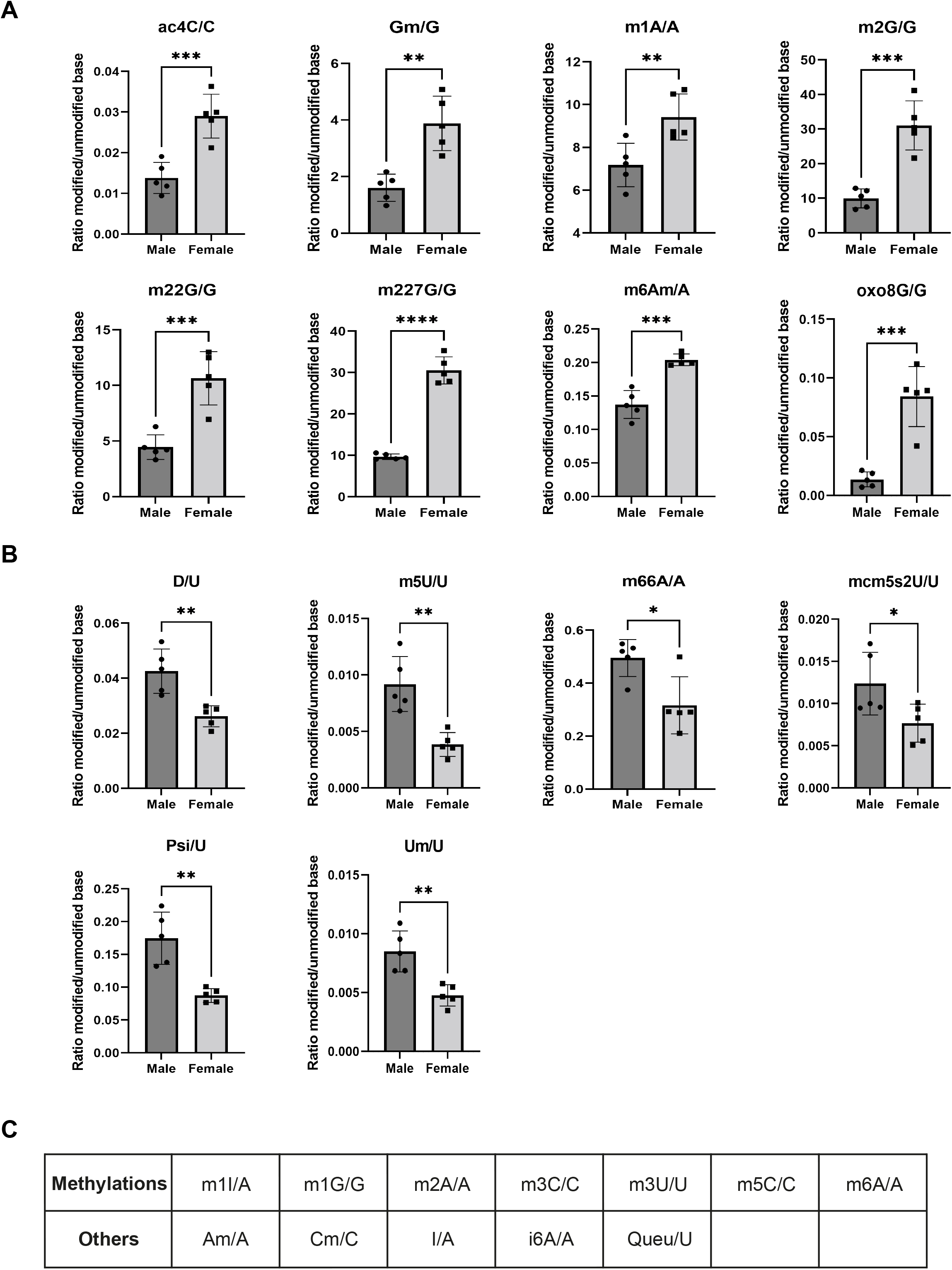
Identification of the modified free nucleosides in the tear film of male and female mice. (**A**) Eight modified free nucleosides were detected predominantly in the tear film from female mice. (**B**) Six modified free nucleosides were detected predominantly in the tear film from male mice. (C) Among the 12 modified free nucleosides which did not present any sexual bias, seven were specific methylations. Data are represented as mean ±SEM, with n=5 per condition. Statistical analysis using Ordinary one-way ANOVA with *, p□<□0.05; **, p□<□0.01; ***, p□<□0.001; ****, p□<□0.0001.

Building on this signal, we profiled the tear-film proteome across three hormonal contexts, i.e. Male, Female, and testosterone-treated Female (Female+T), in 12-week-old mice to test whether androgen exposure remodels lacrimal protein composition. Tear collection rate was comparable across groups (≈0.8 *µ*L/min; **Fig. 2A**), indicating no hormone-dependent modulation of volume. By contrast, total protein concentration was significantly lower in Female than in Male and Female+T (**Fig. 2B**). Label-free proteomics revealed marked sex differences in tear composition that were largely normalized in Female+T (**Fig. 2C**). Focusing on proteins annotated as secreted or extracellular, several factors were more abundant in Male than in Female with an adjusted p<0.05 and were also higher in Male than in Female+T. These Male-enriched proteins included lipocalin-2 (Lcn2), kallikrein-1 (Klk1) and kallikrein-1b11 (Klk1b11), complement factor D (Cfd), serine protease inhibitor A3N (Serpina3n), the Kunitz-type inhibitor Spint1, the WAP-domain protein Wfdc12, zinc-α2-glycoprotein (Azgp1), glutathione peroxidase-3 (Gpx3), and inter-alpha-trypsin inhibitor heavy chain H2 (Itih2). For illustration, Lcn2 showed a Female−Male log2 fold change (log2FC) of −2.93, with Female+T−Male also <0. In parallel, a second set of secreted proteins was more abundant in Female than in Male with adjusted p<0.05 and was also higher in Female than in Female+T. These Female-enriched factors comprised pancreatic lipase-related protein 1 (Pnliprp1), the proprotein convertase Furin, Ovostatin homolog (Ovos), mucin-like protein 2 (Mucl2), and exocrine gland-secreted peptide 1 (Esp1). For example, Pnliprp1 displayed log2FC(Female−Male)=9.40 and log2FC(Female−Female+T)=6.63. Together, these datasets define sex-associated, secretion-focused candidates and document that values in Female+T follow the directions indicated by the corresponding pairwise contrasts.

**Figure 2.**
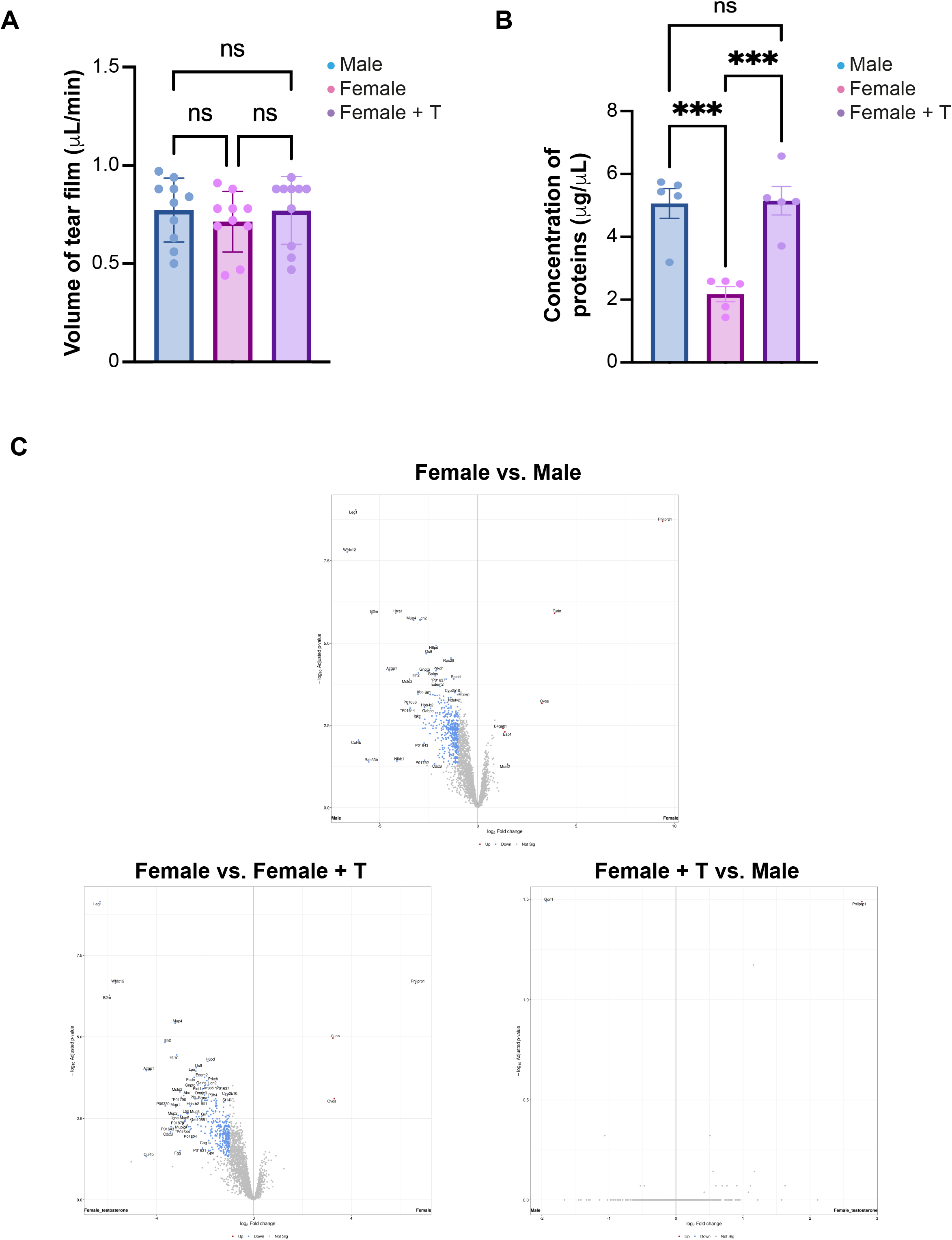
Evaluation of the tear film quantity and quality. (**A**) The tear film is collected and its volume is measured. No significant differences are detected in the tear film secretion when comparing samples from males, females and testosterone-treated females (female + T). (**B**) The protein concentration is significantly higher in testosterone-driven hormonal contexts. (**C**) Volcano plot representations of tears proteomics analysis by mass spectrometry (n=5 per group). For proteomics volcano plots, all identified proteins are represented and the limit of statistical significance is shown with the horizontal dotted line. Genes and proteins with a fold change lower than 0.5 are colored in blue. Genes and proteins with a fold change higher than 2 are colored in red. The genes and proteins with an intermediate fold change are colored in grey. Data are represented as mean ±SEM, with n=5 per condition. Statistical significance was assessed by unpaired two-tailed t-test (ns, non-significant; ***, p□<□0.001).

### Corneal innervation is sexually dimorphic and androgen-responsive

Having mapped the tear-film milieu, we next examined the second pillar of corneal physiology—the dense intraepithelial sensory plexus. Loss of corneal innervation precipitates neurotrophic keratopathy, with progressive epithelial breakdown that can culminate in ulceration and transplantation (Dua *et al*, 2018).

We visualized corneal nerves by βIII-tubulin immunolabeling, a pan-neuronal marker (Meneux *et al*, 2024) (**Fig. 3A**). The neural lattice in males was visibly sparser than in females. Strikingly, androgen exposure shifted females toward a male-like configuration: testosterone-supplemented females displayed similarly thinned meshes. These impressions were borne out by quantitative volumetry. Using our validated fiber-volume pipeline to analyze the central 50% of each cornea (Meneux *et al*, 2024, 2025), female epithelia contained ∼30% more neural volume than either males or testosterone-supplemented females (**Fig. 3B**).

**Figure 3.**
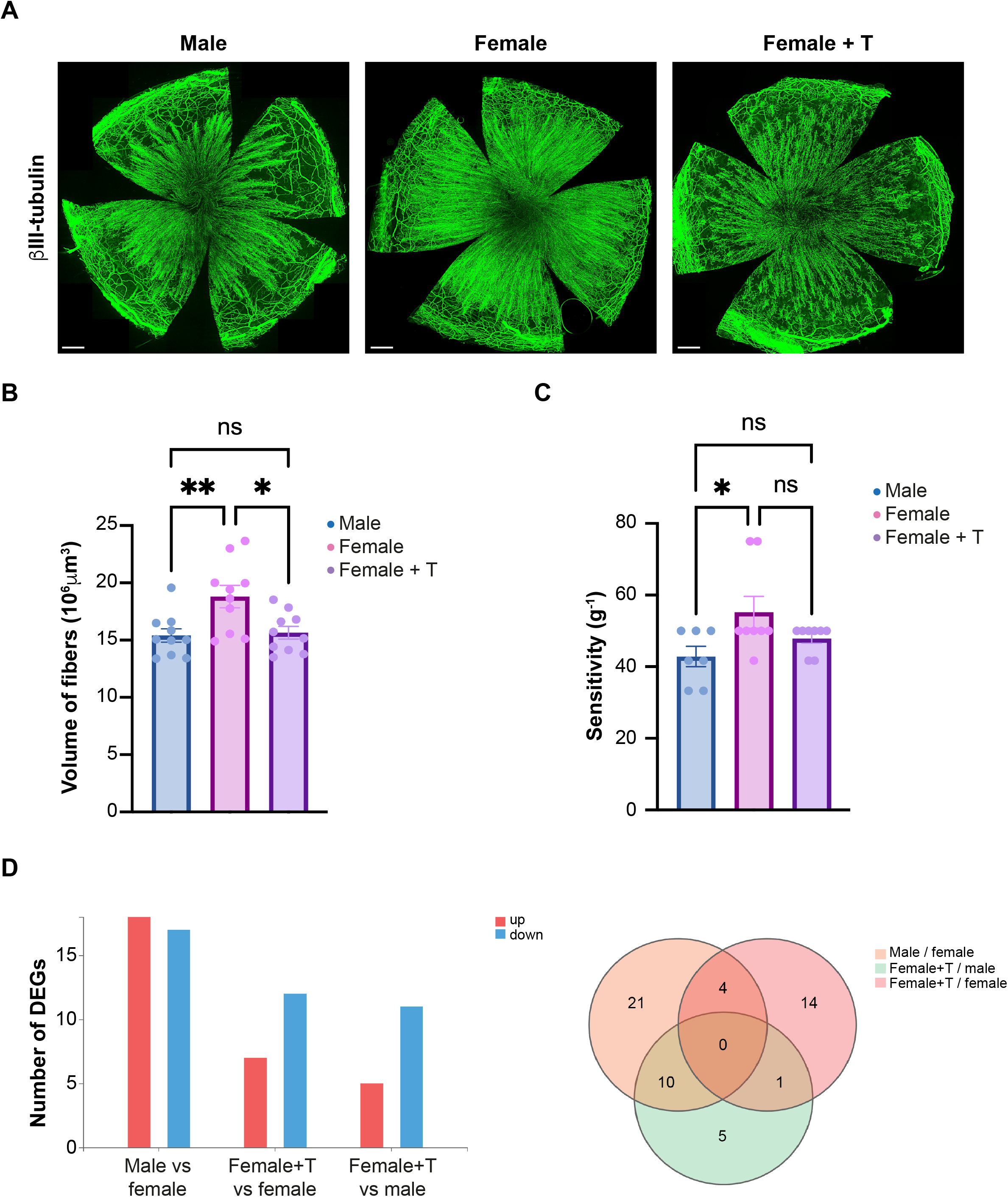
Structural, molecular and functional characteristics of corneal innervation. (**A**) The visualization of βIII-tubulin+ axons in 12 weeks of age male, female and testosterone-treated females (female +T) shows a dense innervation, with a denser pattern in intact females. (**B**) The epithelial nerve fiber volumes in the central area covering 50% of the whole corneal surface are then measured for each stage using Imaris software. Volumes of βIII-tubulin+ fibers are significantly higher in females than in males and testosterone-treated females. (**C**) Central corneal sensitivity is monitored up with Von Frey filaments. The sensitivity decreases in androgene-driven contexts. (**D**) The number of differentially expressed genes (DEG) was the highest between males and females (35 genes), while only 19 DEGs were identified between males and female + T. Scale bars = 300µm. Data are represented as mean ±SEM, with n=6 to 10 per condition. Statistical analysis using Ordinary one-way ANOVA with ns, non-significant; *, p□<□0.05; **, p□<□0.01.

To link structure to function, we measured corneal mechanosensation by von Frey aesthesiometry as previously described (Meneux *et al*, 2024, 2025). Touch thresholds paralleled nerve density: physiological females were significantly more sensitive than both comparison groups, whereas androgenized females were indistinguishable from males (**Fig. 3C**).

To probe the origin of this dimorphism and its modulation by testosterone, we profiled the trigeminal ganglia—the source of corneal afferents (**Fig. 3D**). Thirty-five genes were differentially expressed between males and females. By contrast, only sixteen genes differed between males and testosterone-supplemented females; ten of these overlapped the sex-dimorphic set, indicating partial normalization toward a male-like transcriptional state under androgen exposure. Finally, fourteen were unique to testosterone treatment.

Gene-level examples illustrate three response classes. First, some transcripts were fully androgen-responsive: for instance, *Lcn2* showed a 4.86-fold higher expression in males versus females, which was effectively modified in testosterone-exposed females (female+T vs. male, fold change 0.97). Second, other transcripts remained partially divergent despite treatment: *Fhsb* exhibited a large male–female difference (744.43-fold), not achievable with testosterone exposure (female+T vs. female, 45.89-fold). Third, some genes were unresponsive to testosterone: Y-linked genes (Eif2s3y, Uty, Uba1y) were, as expected, not detected in females (with or without testosterone), and *Katnal1*, a regulator of adult neurogenesis (Lombino *et al*, 2019), was more highly expressed in females and unchanged by testosterone. Notably, expression of fiber-identity markers (*TRPM8, SP, VIP, L1CAM*) (Meneux *et al*, 2025) was unaffected by hormonal state, suggesting that androgens primarily remodel axonal abundance or arborization rather than switching fiber subtype identity.

Together, these findings reveal a previously unrecognized, hormone-tunable sexual dimorphism in corneal innervation, spanning anatomy, sensation, and upstream sensory-ganglion transcription, in which androgen exposure aligns female corneas toward a male-like state.

### Epithelial renewal and molecular landscap are hormonally tuned and spatially re-patterned

Building on the androgen sensitivity of tear chemistry and nerve architecture, we next examined the third pillar of corneal homeostasis—epithelial physiology. In mice, short-term renewal is sustained by Krt14^+^ progenitors distributed across the basal epithelial layer (Kalha *et al*, 2018). To visualize their output, we crossed Krt14-CreERT mice with the multicolor CAG-Cytbow reporter (Loulier *et al*, 2014) and tracked clonal expansion over three weeks across three hormonal contexts. Most fluorescent clones remained punctate, consistent with local replacement (**Fig. 4A**). However, clone density diverged sharply by hormonal milieu: females exhibited the greatest number of labeled foci, whereas both males and testosterone-supplemented females showed markedly fewer events.

**Figure 4.**
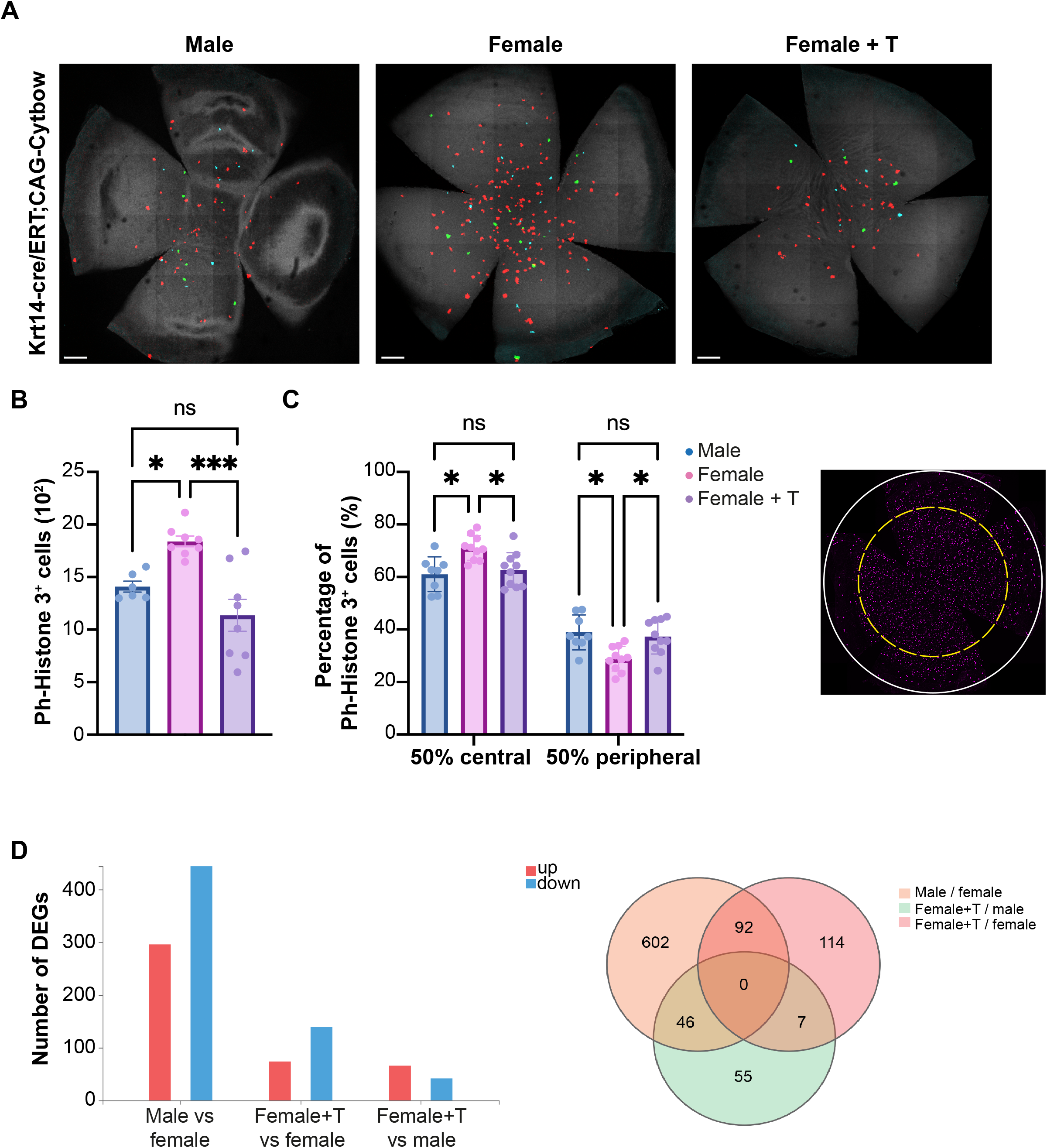
Epithelial corneal renewal dynamics and transcriptomic signature. (**A**) Three weeks after Tamoxifen treatment of Krt14-Cre/ERT;CAG-Cytbow, the fluorescent clones are visible on the whole corneal surface in male, female and testosterone-treated females (female + T), with a higher clone density in females. (**B**) The count of phospho-Histone3^+^ (Ph-Histone3) cells shows a higher proliferation in female corneal epithelium compared to males and Female + T. (**C**) The spatial spreading of these cells displayed a different repartition in females. The illustration represents the limits of the 50% of corneal surface (yellow dotted circle, diameter d/√2) compared to the whole surface (white circle, diameter d). (**D**) The RNA sequencing revealed a strong sexual dimorphism (740 DEGs between males and females), which was mainly lost after testosterone exposure (109 DEGs between males and females + T). Scale bars are 400µm. Data are represented as mean ± SD, with n=8 to 10 per condition. Statistical significance was assessed by Ordinary one-way ANOVA, and two-way ANOVA, with ns, non-significant; *, p□<□0.05; ***, p□<□0.001.

Mitotic activity, quantified by phospho-Histone H3^+^ nuclei in the basal layer, recapitulated the clonal trends—female corneas harbored significantly more proliferating cells than either comparison group (**Fig. 4B**). Spatial mapping revealed a conserved central predominance of proliferation across cohorts, but androgen exposure redistributed renewal: physiological females devoted a higher fraction of mitoses to the central 50% of the cornea, whereas males and androgenized females shifted activity toward the periphery (**Fig. 4C**).

To define the molecular underpinnings of these shifts, we profiled epithelial transcriptomes under the three hormonal conditions. We identified 740 differentially expressed genes (DEGs) between male and female corneas, underscoring a substantial sexual dimorphism (**Fig. 4D**). Testosterone treatment in females altered 213 genes, 43% (92/213) of which overlapped the sex-dimorphic set. Consistently, androgenized female corneas converged toward the male transcriptional state, with only 105 DEGs distinguishing those two contexts.

Mirroring the trigeminal ganglion analysis, the transcriptomic comparisons revealed three gene-response classes. First, sex-dimorphic transcripts included Tgoln2, which was enriched in females, and Cf5, which was ∼6-fold higher in males versus females and ∼3.43-fold higher in testosterone-treated females versus untreated females, indicating partial androgen-driven convergence. Second, strongly androgen-responsive genes were exemplified by Muc5b, which increased ∼94.35-fold in testosterone-treated females relative to females. Third, testosterone-unresponsive genes included Nectadrin (Cd24a), whose expression was lower in males than in females (fold change 0.70, male vs. female) and remained unchanged between untreated and testosterone-treated females.

Together, these data establish that androgen signaling tunes corneal epithelial renewal in vivo, dampening progenitor output, redistributing proliferation spatially, and reprogramming the epithelial transcriptome toward a male-like state, revealing a previously unrecognized, hormone-responsive layer of corneal homeostasis.

## Discussion

This study reveals a coordinated sexual dimorphism across the tear film, epithelial renewal, and sensory innervation, and indicates that each pillar is plastically tuned by circulating androgens rather than fixed by development. In the tear film, the overall proteomic landscape differed markedly between sexes yet shifted toward the male state when females received testosterone, consistent with a hormone-responsive signature. Within the secreted proteome, the patterns point to concerted changes in lipid transport and processing, protease–antiprotease balance, complement activity, and secretory/mucin-associated pathways—axes that align with downstream differences observed in corneal epithelial dynamics and sensory function. The reduction in female– male divergence under testosterone further supports a reversible remodeling of lacrimal outputs, linking endocrine state to surface homeostasis.

These findings have practical implications. First, they argue against pooling male and female samples in discovery studies and support sex-stratified reference ranges for tear-based diagnostics. Second, they suggest that hormonal status can confound or modulate ocular surface biomarkers, with relevance for conditions such as dry eye, neuropathic pain, and neurotrophic risk. Finally, by coupling a sex-stratified tear metabolite profile with a hormone-sensitive proteome, the work frames a tractable path to mechanism-guided biomarkers and interventions that account for endocrine context at the ocular surface.

The neural analyses extend this molecular dimorphism to structure and function. Female corneas displayed denser intraepithelial nerve meshes and lower touch thresholds than males, and exogenous testosterone reduced both fiber volume and mechanosensory sensitivity in females toward the male state. Transcriptomic profiling of trigeminal ganglia—the source of corneal afferents—identified sex-biased expression in pathways related to axon guidance, neuropeptide processing, and mitochondrial energetics, offering plausible molecular substrates for the observed anatomical and sensory differences. Given the clinical use of trophic agents for neurotrophic keratopathy, these data raise the possibility that dose, timing, and expected effect size may need to be sex-specific, and that peri-therapeutic hormonal status could modulate outcomes.

Epithelial physiology showed a similarly coherent pattern. Clone tracking from Krt14^+^ basal progenitors and phospho-histone H3 counts indicated that females cycle faster and produce more labeled foci than males, whereas androgenized females reverted toward the male state. Spatial mapping revealed a conserved central predominance of proliferation across cohorts but also showed that androgens redistribute renewal toward the periphery. Bulk RNA-seq substantiated a large sexual dimorphism and showed that testosterone shifts the female transcriptome toward the male state, implicating pathways such as Wnt, Hippo, and extracellular-matrix signaling that are known to govern stem-cell behavior. Considering the central role of limbal epithelial stem cells in corneal maintenance and repair, these findings suggest that circulating hormones modulate both the rate and the topology of epithelial renewal, with potential implications for wound coverage, barrier integrity, and graft integration.

Viewed together, these datasets converge on androgen signaling as a coordinating node that shapes the chemical environment of the tear film, remodels sensory architecture and mechanosensation, and tunes progenitor output and the spatial blueprint of epithelial proliferation. The rapidity and reversibility of the changes we observe with exogenous testosterone argue for dynamic hormonal control over maintenance programs rather than static sexual fate. Clinically, this framework supports several testable propositions: sex-stratified reference intervals could improve the interpretability of tear-based biomarkers; hormone-aware interventions—topical or systemic— might be deployed to dampen hyper-innervation–linked pain or to boost epithelial regeneration after injury or surgery; and sex matching or perioperative hormonal modulation may merit evaluation as variables in corneal transplantation. Trial design for lubricants, neuroprotectants, and regenerative biologics should incorporate sex as a biological variable from inception, with pre-specified subgroup analyses.

In sum, the cornea is not a passive window but an active sensor–effector organ whose maintenance programs are deeply sex-tuned and hormonally plastic. Recognizing and leveraging this dimorphism sharpens mechanistic understanding and opens therapeutic avenues, from biomarker design to individualized regeneration strategies. As precision medicine advances, ocular health must keep pace; the present work lays a foundation for sex-aware diagnostics and interventions at the ocular surface.

## Material and Methods

### Animals included in this study

Mice were housed in a 12 h light/12 h dark cycle with free access to food, and animal procedures were carried out in accordance with institutional, and ARRIVE guidelines. Animal protocols were approved by Languedoc Roussillon animal experimentation ethics committee (CEEACD/N°36), and the Ministère de la Recherche et de l’enseignement Supérieur (authorization 2016080510211993 version2). All of the procedures were carried out in accordance with the French regulation for the animal procedure (French decree 2013-118) and with specific European Union guidelines for the protection of animal welfare (Directive 2010/63/EU). Swiss/CD1 female and male mice (RjOrl:SWISS, Janvier Labs, France) were used for most experiments.

### Hormonal treatment

Female mice receiving the testosterone treatment were 6 weeks of age at the time of their first injections. Mice under isoflurane anaesthesia received twice-weekly mid-back 100-μL subcutaneous injections of Testosterone (T) enanthate in sesame oil. T enanthate at 0.45 mg per dose was diluted from the stock solution (200 mg/mL, CarlRoth, dissolved in sesame oil, Sigma). After 6 weeks of T injections, corneal sensitivity and tear sampling were performed on all mice. Then, mice were euthanized and eyes and trigeminal ganglions harvested for histology and omic analyses.

### Assessement of corneal sensitivity

Sensitivity was measured using Von Frey filaments (Bioseb, France, reference bio-VF-M). These filaments, applied to the center of the cornea of an immobilized mouse, exert a force ranging from 0.008 to 0.6 g. Filaments of increasing stiffness were used until an eye-blink reflex was elicited. The test was performed over three consecutive days, and the data collected for each cornea were averaged. Sensitivity was expressed in g^−1^. All procedures were carried out by the same experimenter under single-blind conditions.

### Tear film collection

Tear samples were collected from conscious, hand-restrained mice using disposable 1 *µ*L glass microcapillary tubes (reference 9000101, Hirschmann). The microcapillary was gently applied to the outer corner of each eye, avoiding direct contact with the ocular surface, for one minute per eye, with approximately one tap per second. Samples from both eyes were then combined into a single tube per mouse in 12 *µ*L of ammonium bicarbonate (ABC) buffer provided by the proteomics platform. The samples were kept on ice at 4 °C throughout the collection process, then stored at −80 °C for long-term preservation.

### Proteomic analysis

The proteomic profile of tear samples was analyzed. Total protein concentration was measured using a NanoDrop One Microvolume UV–Vis Spectrophotometer (Thermo Fisher Scientific, reference ND-ONE-W). For each sample, 3□*µ*g of total protein were reduced, alkylated, and digested with trypsin using the automated SP3 protocol on the Bravo 96LT AssayMap Protein Sample Prep Platform (Agilent Technologies). Following digestion, peptides were desalted and injected in technical triplicates on the Evosep One system coupled to a timsTOF HT mass spectrometer (Bruker Daltonics). Protein identification was performed using DIA-NN software (version 1.8.1) with the following parameters: trypsin as the digestion enzyme, allowing one missed cleavage; minimum peptide length of seven amino acids; precursor charge range of 1 to 4; precursor m/z range set between 300 and 1800; and a protein-level false discovery rate (FDR) threshold of 1%. The mouse proteome from the UniProt database (Release_2024_03) was used as the reference. Sample preparation-induced modifications were accounted for, with asparagine deamidation and methionine oxidation included as variable modifications, and cysteine carbamidomethylation as a fixed modification. A maximum of two variable modifications per peptide was allowed.

Label-free quantification (LFQ) intensities were processed using Perseus software (version 1.6.15.0) and the DIA-Analyst platform (version 0.8.6). The average intensity of the three technical replicates was calculated for each protein and used for downstream statistical analysis. LFQ values were log_2_-transformed and grouped by experimental condition (male, female, and testosterone-treated female) for each eye (left and right). Proteins with complete data (100% valid values) across biological replicates were retained for analysis. Significantly regulated proteins in each pairwise comparison of matched samples were identified using an adjusted p-value threshold of 0.05 (t-statistic correction) combined with a log_2_(fold change) cutoff of ±1.

### Tissue collection and processing

For each sample, the two trigeminal ganglions (right and left) from a same animal were collected and pooled in a single tube, snap frozen then stored at -80°C (n = 5 mice). Eyes were collected using curved scissors to cut the optic nerve. Corneas and trigeminal ganglions for transcriptomic analysis were directly dissected and pooled (right and left) in a single tube snap frozen then stored at -80°C (n = 5 mice). For immunofluorescence labeling, the eyes were fixed in 4% PFA (Antigenfix) for 20 minutes, then washed in PBS. The eyes were then dehydrated for 2H in 50% ethanol/PBS and stored in 70% ethanol/PBS at 4°C.

### RNA isolation and cDNA library synthesis

Frozen cornea and trigeminal ganglion tissues were shipped on dry ice to BGI (Shenzhen, China) for library preparation and sequencing on the DNBseq platform. Total RNA was extracted with the RNeasy Mini kit (Qiagen). Poly(A)+ mRNA was enriched using oligo(dT)-conjugated magnetic beads and subsequently fragmented. RNA integrity and fragment size distributions were verified on a Fragment Analyzer, and only samples meeting quality criteria proceeded to library construction. Libraries were prepared with the BGI Optimal Dual-mode mRNA Library Prep Kit and sequenced on a DNBSEQ-T7 instrument (paired-end 2×100 bp; ∼50 million clean reads per library). Raw reads were processed with SOAPnuke (Li *et al*, 2008) to remove adapters, contaminants, and low-quality sequences. Clean reads were aligned to the mouse reference genome (*Mus_musculus_10090; UCSC mm10 v2201*) using HISAT2, and gene-level quantification was performed with HTSeq-count to generate the raw count matrix across all samples.

### Transcriptomic data analysis

These raw counts were analysed using Dr. Tom multi-omics data mining system (https://biosys.bgi.com). In short, Bowtie2 (Langmead & Salzberg, 2012) was applied to align the clean reads to the gene set, in which known and novel, coding and noncoding transcripts were included. Expression levels of gene were calculated by RSEM (v1.3.1) (Li & Dewey, 2011). The heatmap was drawn by pheatmap (v1.0.12) according to the gene expression difference in different samples. Essentially, differential expression analysis was performed using the DESeq2(v1.34.0) (Love *et al*, 2014) with Q value ≤ 0.05 (or FDR ≤ 0.001). Finally, GO (www.geneontology.org) and KEGG (www.kegg.jp) enrichment analysis of annotated different expression gene was performed by Phyper based on Hypergeometric test. The significant levels of terms and pathways were corrected by Q value with a rigorous threshold (Q value ≤ 0.05).

### Immunofluorescence labeling on whole cornea

Previously collected eyes were rehydrated in 50% ethanol for two hours and rinsed in PBS. Corneas were dissected just above the ciliary body, along the limbal region using microdissection scissors. They were then immersed in blocking and permeabilization solution in FSG 5% (Sigma-Aldrich, reference G7765) and goat serum (GS) 5% (Thermo Fisher Scientific, reference 16,210,064) in 0,5% PBS/Triton X-100 at room temperature. Then the corneas were incubated overnight at 4°C with primary antibodies and then secondary antibodies (**Table x**). After washing, nuclei were stained with Hoechst (Thermo Fisher Scientific, reference H3570) for 10 minutes. The corneas were incised at four cardinal points using a 15C carbon steel surgical blade (reference 0221, Swann-Morton) and then flat-mounted in Vectashield medium (Vector Laboratories, H-1000), with the epithelium positioned against the coverslip.

### Images acquisition and processing

The acquisition of the images was performed as previously described (Meneux *et al*, 2024). A Leica Thunder Imager Tissue microscope was used to acquire the whole-cornea images, using the navigator module with the large volume computational clearing (LVCC) process. The LAS X software (version 3.7.4) was used to obtain the images using a 20X/0.55 objective. Then, to process the images Imaris Bitplane software (version 10.1) was used. All of the images from a single panel were acquired and processed with the same parameters.

### Innervation volume measures and cell counting

A centered circle covering 50% of the corneal area was drawn using the Fiji measurement plug-in (FIJI (RRID:SCR_002285)). The corneal diameter was measured twice for each sample, and the mean value was used to determine the circle’s radius. A 50% circular crop of each cornea was then generated using the Crop plug-in, followed by the Clear Outside plug-in. The cropped images were converted to .ims format using the Imaris converter. Innervation density was quantified using the surface tool in Imaris. Once the surface was defined, volume data (expressed in μm^3^) were extracted for statistical analysis. Proliferating cells were counted on Fiji using the binary convert to mask plug-in and analyze particles plug-in.

### Genetic cell lineage tracing

Transgenic mice broadly expressing CAG-Cytbow transgene (Loulier *et al*, 2014) were modified to achieve random expression of mTurquoise2, mEYFP, tdTomato or mCherry from the broadly active CAG promoter following Cre recombination (Abdeladim *et al*, 2019; Clavreul *et al*, 2019). CAG-Cytbow mice were crossed with Krt14-cre/ERT animals (The Jackson Laboratory), and their Krt14-cre/ERT;CAG-Cytbow offspring analyzed 3 weeks after three subcutaneous injections of Tamoxifen (200 μL at 20mg/ml; Sigma, T5648) dissolved in corn oil (Sigma, T5648) at 9 weeks of age. We collected the eyes at 12 weeks of age, fixed them in 4% PFA (Antigenfix) for 20 minutes, then washed them in PBS. The corneas were dissected and mounted in Vextashield Plus (Vector Laboratories), and then stained with bio Tracker NIR694 nuclear Dye (Sigma-Aldrich, SCT117, 1/5000). A confocal microscope Zeiss LSM880 airyscan was used to acquire the whole cornea images.

### Statistical analysis

Data analysis was performed using GraphPad Prism software (version 10.2.2). Results are presented as mean ± SEM. Significant p-values were represented as *p < 0.05, ** p < 0.01, *** p < 0.001, **** p < 0.0001. Statistical analyses comparing mean values were performed using one of the following tests: the Friedman test with a Dunn’s multiple comparison test, the Kruskal-Wallis test with a Dunn’s multiple comparison test, or a Ordinary one-way ANOVA followed by Dunnett’s multiple comparison test. Simple comparisons were performed using an unpaired t-test. All tests were identified in the legend of each figure.

## Data availability

The raw data is available upon request and the datasets produced for the current study are publicly available. The RNA-Sequencing data is available in the GEO repository, under the accession number xxx. The proteomic data is available in the PRIDE repository, via ProteomeXchange, with the identifier xxx.

## Author contributions

Conceptualization: N.F., A.C.M, K.L., C.H., F.M.; Methodology: N.F., A.C.M., S.P., M.G., M.D., L.F., A.A., J.V., A.D., V.D., K.L., C.H., F.M.; Validation: J.V., A.D., V.D., K.L., C.H., F.M.; Formal analysis: N.F., A.C.M., S.P., M.G., M.D., A.A., J.V., K.L., C.H., F.M.; Data analysis: N.F., A.C.M., S.P., M.G., M.D., L.F., A.A., J.V., K.L., C.H., F.M.; Writing: N.F., A.C.M., S.P., M.G., M.D., L.F., A.A., J.V., A.D., V.D., K.L., C.H., F.M.; Supervision: F.M.; Project administration: F.M.; Funding acquisition: F.M.

## Acknowledgments

This research was supported by ATIP-Avenir program, Inserm, the Région Occitanie, ANR (ANR-21-CE17-0061, TeFiCoPa), FRM (REP202110014140), Support for research: I-SITE 2024 - program of excellence of the University of Montpellier, CBS2 Doctoral School, the Fondation Groupama. Mass spectrometry and epitranscriptomic experiments were carried out using the facilities of the Montpellier Proteomics Platform (PPM, BioCampus Montpellier), a member of the national Proteomics French Infrastructure (ProFI UAR 2048) supported by the French National Research Agency (ANR-24-INBS-0015, Investments for the future F2030). We thank the MRI-DBS imaging facility, member of the national infrastructure France-BioImaging (https://ror.org/01y7vt929) supported by the French National Research Agency (ANR-24-INBS-0005 FBI BIOGEN). We thank the personel of the RAM-INM animal core facility, member of the Animal Facility Network in Montpellier (RAM).

